# Recapitulation of clinical and molecular hallmarks of lipid-induced hepatic insulin resistance in a zonated, vascularized human liver acinus microphysiological system during metabolic dysfunction-associated steatotic liver disease (MASLD) progression

**DOI:** 10.1101/2025.11.02.686163

**Authors:** Julio Aleman, Lawrence Vernetti, Mark E. Schurdak, Richard DeBiasio, Greg LaRocca, Vijay K Yechoor, D. Lansing Taylor, Andrew M. Stern, Mark T. Miedel

**Affiliations:** Organ Pathobiology and Therapeutics Institute; University of Pittsburgh Department of Bioengineering; University of Pittsburgh Department of Computational and Systems Biology; University of Pittsburgh Liver Research Center; University of Pittsburgh Diabetes and Beta Cell Biology Center, Division of Endocrinology and Metabolism; University of Pittsburgh Department of Pharmacology and Chemical Biology

**Keywords:** Metabolic dysfunction-associated steatotic liver disease (MASLD), Human microphysiology systems (MPS), Metabolic disease mechanism, Hepatic steatosis, Hepatic insulin resistance, Hyperglycemia, Dyslipidemia, Liver zonation

## Abstract

Metabolic dysfunction-associated steatotic liver disease (MASLD) impacts ca. 30% of the global population and is very heterogeneous making it a challenge to produce therapeutics. The heterogeneity arises from genetics, co-morbidities, the microbiome and lifestyle. To help address this challenge, we have refined the human vascularized liver acinus microphysiological system (vLAMPS), which provides an all-human platform for drug development, in line with recently updated federal requirements for the use of New Approach Methodologies (NAMs). By introducing clinically relevant media perturbations and employing several diverse and reproducible *in situ* and systemic measurements, we show that the vLAMPS can recapitulate key structural and functional aspects of normal physiology, acinus zonation, and all stages of MASLD progression including stellate cell activation and fibrosis. Importantly, in this study we also demonstrate that several hallmarks of lipid-induced hepatic insulin resistance paralleled MASLD progression. These included diminution of insulin receptor substrate 2 (IRS2) protein, compromised insulin receptor mediated insulin clearance, enhanced pericentral lipid accumulation, increased VLDL secretion, and enhanced hepatic glucose output mediated by increased periportal nuclear translocation of FOXO1. These results suggest that the mechanisms underlying MASLD progression in vLAMPS are clinically relevant and support the tenable hypothesis that the hepatic insulin resistant state plays both a causal and consequential role in a vicious cycle driving disease progression.

## Introduction

Metabolic dysfunction-associated steatotic liver disease (MASLD) affects approximately 34% of the U.S. population with a similar prevalence among many global populations.^2^ MASLD is the most common cause of chronic liver disease and leading indicator for liver transplant in the US.^3^ Although the prevalence of the progressive form, metabolic dysfunction-associated steatohepatitis (MASH) that increases the risks for cirrhosis, hepatic decompensation and hepatocellular carcinoma is lower (6-8%)^2,4–6^ it has been projected that over the next 25 years the prevalent cases of decompensated cirrhosis will more than triple, the incident cases of liver cancer will double and liver transplants will almost quadruple.^2^ MASLD is closely linked to other chronic metabolic disorders including obesity, type 2 diabetes mellitus (T2DM), chronic kidney disease (CKD), and cardiovascular disease (CVD).^7^ Demonstrating the strong association with these comorbidities, the global prevalence of MASLD is 65% in the subgroup of patients with T2DM with half of these patients having MASH^8^ and CVD is the leading cause of death in persons with MASLD.^3^ Furthermore, MASLD has been shown to be a risk factor for new onset heart failure,^9,10^ atrial fibrillation,^11^ T2DM,^12^ CKD,^13^ and certain extrahepatic cancers.^14^ Together, these results indicate that the liver plays a key role in the pathophysiology of these metabolism-related disorders that constitute the Cardiovascular-Kidney-Metabolic (CKM) syndrome^7^ with MASLD preceding extrahepatic dysfunction in most cases. Identifying targetable hepatic mechanisms that drive the complex pathophysiology of MASLD could help meet the urgent need for effective MASLD therapies and broaden treatment options for the CKM syndrome.^15–17^

Several lines of evidence support lipid-induced hepatic insulin resistance as a major pathogenic mechanism in MASLD.^18–20^ Insulin, synthesized and secreted from the beta cells of the pancreas, promotes glucose storage through glycogenesis, stimulates *de novo* lipogenesis (DNL) to convert excess carbohydrates to lipids as a storable source of energy, and regulates the nuclear translocation of the transcription factor Forkhead box protein O1 (FOXO1) to suppress gluconeogenesis and glycogenolysis.^21,22^ In the insulin resistant state (IR), hepatic uptake of glucose and insulin clearance are compromised while hepatic glucose synthesis and DNL are increased promoting a loss of metabolic homeostasis. The increase in DNL promotes excess triglyceride and very low density lipids (VLDL) formation and secretion accompanied by lipid-induced hepatocyte inflammation and injury.^23,24^ A vicious cycle ensues causing release of proinflammatory cytokines and damage-associated molecular patterns (DAMPs) that promote stellate cell activation leading to fibrosis and associated cirrhosis and hepatocellular carcinoma (HCC).^25,26^

Several elegant studies in rodent models have identified many of the molecular mechanisms involved in lipid-induced hepatic insulin resistance.^27–30^ However, several aspects of MASLD progression have proven challenging to recapitulate in mouse models making it difficult to determine the precise role of lipid-induced insulin resistance in the pathogenesis of MASLD.^31^ Furthermore, the molecular basis for how DNL is dysregulated in the context of hepatic IR, a key question as discussed above with important therapeutic implications, remains controversial requiring further investigation^32^. Studies in patients have corroborated some of the results from the rodent models,^33,34^ but an enhanced analysis requiring a comprehensive set of measurements that are limited in the clinical setting is still required to deconstruct lipid-induced hepatic insulin resistance mechanisms to understand their role in MASLD pathogenesis.

To address species differences and enable use of both primary human and patient-derived iPSC liver cells, multiple laboratories have developed all-human, multi-cellular liver microphysiological systems (MPS) that replicate key structural and functional features of the hepatic acinus and serve as a reference for MPS constructed with iPSC-derived liver cells.^1,35–38^ In the studies here, we have implemented the vascularized liver acinus microphysiological system (vLAMPS), a two-chamber device with a flexible middle section composed of a porous membrane that enables tissue culture in both the apical and basolateral sides (Figure 1). The biomimetic structure of the vLAMPS replicates hepatic sinusoidal periphery, extracellular matrix composition, and parenchyma metabolic zonation through the middle section. The device is assembled with four key human liver cell types (hepatocytes, liver sinusoidal endothelial, Kupffer cells, and hepatic stellate cells). The structural arrangement of the four cell types facilitates heterotypic cell communication. This is accomplished by mimicking the space of Disse and enhancing hepatocyte polarization through a liver-specific extracellular matrix.^39^ Additionally, the glass microfluidic device facilitates control over the passive oxygen diffusion, flow rate, and cell metabolism to facilitate the generation of a continuous metabolic oxygen zonation in similar magnitude to the oxygen concentrations of the liver acinus (Figure 1). The oxygen tensions of the *in vivo* liver acinus goes through a 50% decrease, starting at the periportal blood range of 84-91 µmol L^-1^ and decreasing to 42-49 µmol L^-1^ at the pericentral blood, with an intermediate value for the midzonal zone.^40,41^ In the vLAMPS we have reported a similar ∼50% decrease in the oxygen tension at the hepatocytes level, such gradient ranges from of ∼120 to 65 µmol L^-1^ through the parallel perfusion of a glass device.^1^ This device has minimum nonspecific binding of drugs encompassing diverse structures or biologics and permits real-time imaging monitoring.^35^ Integrating *in situ* monitoring capabilities with enhanced efflux screening of multicellular systems provides the first step toward next-generation high-content data analysis.

**Figure 1.**
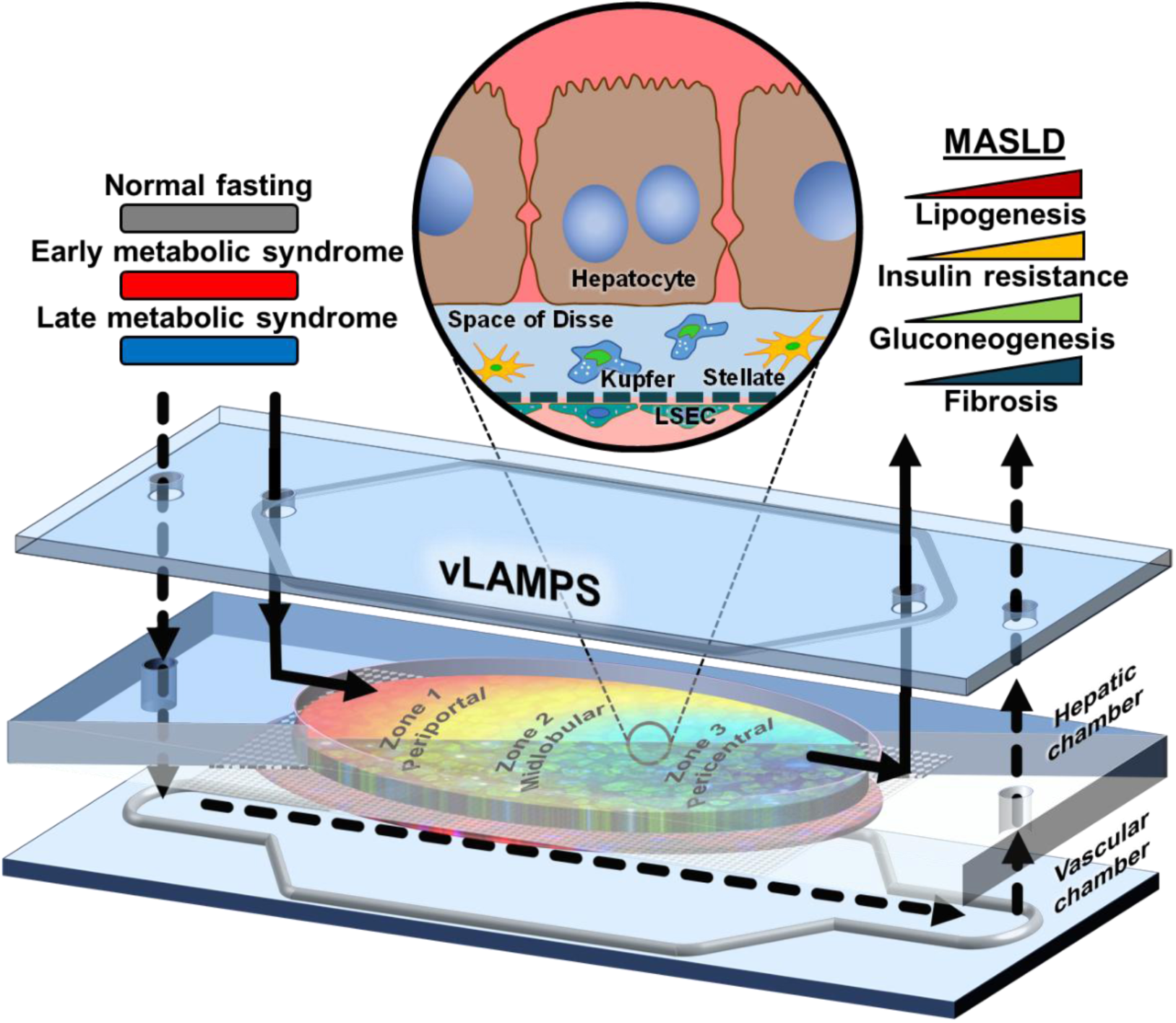
vLAMPS design and influx media composition driven MASLD progression. The vLMAPS is assembled in a double chamber configuration, separated by a porous mebrane. The basolateral side of the membrane conforms the vascular chamber and the apical side the hepatic chamber. Each chamber accomodates their own inlet (downward black arrows) and outlet (upward black arrows), from which one of the three mediums (NF-grey, EMS-red, LMS-blue bars) is perfused in parallel. Thus, the hepatic chamber allows for sampling corresponding to the liver’s interstitial fluid, and the vascular chamber to the sinusoidal fluid. From the collected medium, functional assays can be carried out to determine clinically relevant readouts of lipids, insulin, glucose, and fibrosis macromolecules. Throughtout the membrane (expanded black circle), the hepatic chamber is populated with primary human hepatocytes and the non-parenchymal cells (NPCs) stellate and Kupffer-like cells, mirroring the hepatic interstitial space. The vascular chamber is lined with fenestrated liver sinusoidal endothelial cells, allowing unique permeability between sinusoid and parenchyma. By bringing together the cellular configuration and parallel perfusion into the glass-based device, the vLAMPS replicates the acinus’ metabolic zonation. Starting with the periportal-zone 1, with a oxygen tension ranges between 112.5±6.5 µmol L^-1^. Immediately after, the oxygen diffuses into the parenchyma and is consumed by the cells of the hepatic chamber. As flow progresses, the available oxygen decreases to 89.5±5.5 µmol L^-1^, the midlobular area (zone 2). Lastly, the pericentral zone (Zone 3) reaches oxygen levels of 68±3 µmol L^-1^, representing the lowest yet metabolically functional concentrations.^1^

Utilizing previously established media formulations that mimic the progression of MASLD by modulating key media components including glucose, insulin, free fatty acids and profibrotic drivers including transforming growth factor TGFβ1and LPS^1,42,43^, basal metabolic functions of the liver have been replicated when perfusing normal fasting (NF) medium to generate the vLAMPS continuous oxygen gradient.^1^ Introduction of elevated levels of glucose, insulin, and free fatty acids that recapitulate the levels observed in patients with early metabolic syndrome and defined as EMS medium induced a functional and phenotypical profile in vLAMPS similar to the early stages of MASLD progression. This profile includes hepatic steatosis and inflammation, the secretion of hepatokines that dysregulate pancreatic function, early indicators of stellate cell activation, and fibrosis.^1^ Herein, we extend the vLAMPS capabilities beyond NF and EMS conditions. The system was exposed to known molecular drivers of MASLD that are delivered through a late metabolic syndrome (LMS) medium.^42,43^ Under these new conditions vLAMPS mimics the latter stages of MASLD progression. Furthermore, we have incorporated several additional liver zonation-dependent measurements. We show through this set of functional and phenotypical readouts that hepatic insulin resistance and MASLD progression are tightly associated supporting the use of vLAMPS as a platform for translating mechanistic and pharmacological studies into the clinical setting.

## Results

The progression of Metabolic Dysfunction-Associated Steatotic Liver Disease (MASLD) follows a characteristic sequence of clinically defined stages, advancing from simple steatosis to steatohepatitis, fibrosis and ultimately to cirrhosis. These stages are accompanied by key cellular, metabolic, and spatiotemporal events.^25,44^ The vLAMPS was treated with clinically relevant and chemically defined media (Supplementary Table 1) to mirror the normal fasting (NF), early metabolic syndrome (EMS), and late metabolic syndrome (LMS) conditions. We then implemented diverse phenotypic and functional readouts (Supplementary Table 2-3) with clinical relevancy to determine the extent of MASLD progression and its underlying mechanisms.

### Characterization of basic liver functions during MASLD progression

The vLAMPS was first evaluated to determine that it could maintain basic liver functionality with low cell damage under the three media formulations (NF, EMS, and LMS) for a period of at least 8 days. The three media formulations maintained basic liver functions with detectable albumin and urea levels across the 8 days (Supplementary Figure 1). Albumin plays a key role in managing excess free fatty acids in the bloodstream.^45,46^ In the vLAMPS, albumin levels were highest under EMS conditions throughout the 8-day period, exceeding those observed in NF and LMS conditions. This finding aligns with clinical observations, with LMS in particular, as patients with advanced fibrosis^47^ and insulin resistance^48^ and insulin resistance^48^ typically exhibit lower albumin levels. Biological mechanisms further support these results, as insulin drives albumin synthesis,^49^ explaining the increased albumin levels observed in the higher insulin-containing EMS medium compared to NF. Conversely, inflammation is known to enhance albumin catabolism,^50,51^ which aligns with the lower albumin levels in the LPS/TGF-β-containing LMS medium when compared to EMS. The levels of released LDH were monitored for cell membrane damage. A spike of LDH release specifically in the interstitial compartment under all three media conditions was evident at 2 days suggestive of a pre-steady state artifact resulting from incomplete adaptation of cells to their microenvironments and to flow. Significant LDH release from the hepatic chamber was observed in EMS and LMS compared to NF starting on day 4 consistent with the expected disease specific effect of hepatocyte injury under EMS and LMS. (Supplementary Figure 1 C).

### Impact of circulating free fatty acids on steatosis and lipogenesis

A characteristic phenotype of the liver acinus is the enhanced *de novo* lipogenesis activity within the pericentral region (zone 3).^52,53^ We have previously reported that in comparison to NF conditions, EMS induces a significant increase in lipid droplet accumulation in all three zones with the highest increase characteristically occurring in zone 3.^1,42^ The addition of proinflammatory components as disease drivers in the LMS medium increases hepatic steatosis in all zones of the liver (Figure 2 A and B), while maintaining the pericentral to periportal gradient.^53^

**Figure 2.**
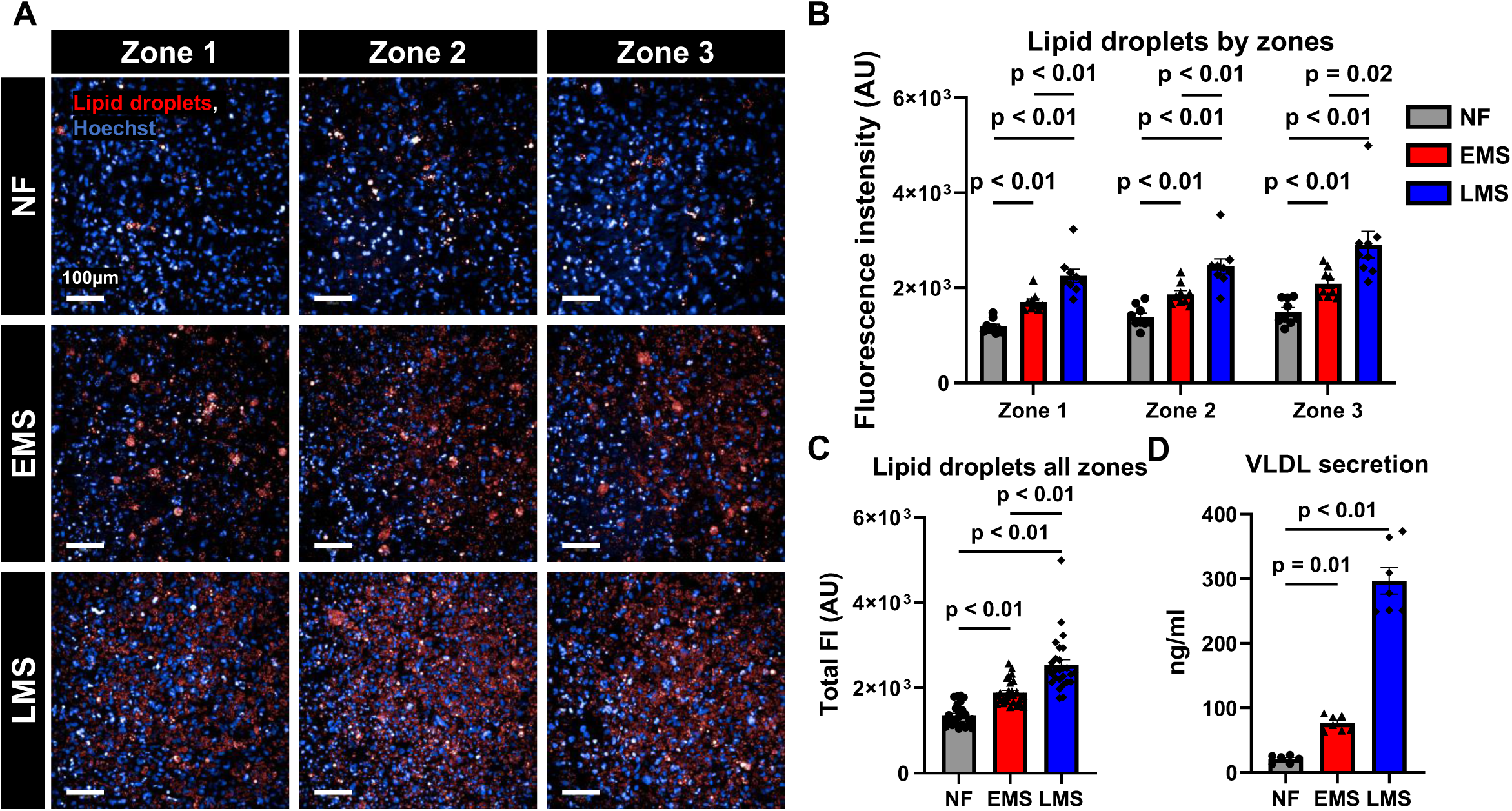
In vLAMPS MASLD severity is associated with hepatic lipid accumulation and VLDL secretion. (**A**) Representative fluorescence images of lipid droplets (LipidTOX) in hepatocytes under NF, EMS, and LMS conditions in their respective zones on day 8. Scale=100µm. (**B**) The quantification by zones of the LipidTOX signal intensity exhibits the zone-specific lipid profile of the vLAMPS. (**C**) The averaged quantification of the LipidTOX signal intensity exhibits significant steatosis in EMS and LMS. Scatter plot graphs display the mean and standard error of mean (SEM) of the fluorescence intensity of the LipidTOX signal of 8 fields at the ROI of each metabolic zone in the vLAMPS, n = 10 MPS. For graph -B-statistical analysis by Two-Way ANOVA with multiple comparisons by Fisher’s LSD test to individually compare media conditions with their respective zone with p-values shown. For graph -C-, statistical analysis by Kruskal-Wallis with multiple comparisons by Dunn’s test was used to compare all three media, with p-values shown. (**D**) VLDL secretion at day 8 at the vLAMPS hepatic chamber efflux. Scatter plot graphs display the mean and standard error of mean (SEM), n = 7 MPS. Graph - D - statistical analysis was performed using One-way ANOVA with multiple comparisons using the Dunnett test to compare EMS and LMS to NF medium with p-values shown.

To avoid potential hepatocyte lipotoxicity caused by their excessive accumulation, these lipids, mostly in the form of triglycerides and cholesterol, are secreted from the hepatocyte as VLDLs.^54^ Their increased secretion is a clinical characteristic of MASLD progression and is mechanistically linked to hepatic insulin resistance.^54^ On day 8 EMS and LMS conditions significantly enhanced the VLDL signal compared to NF (Figure 2 D). LMS induced significantly higher levels than EMS. Thus, it can be inferred that the excess VLDL reflects a hepatocyte protective mechanism, emerging from the excessive intracellular lipid droplets, particularly those induced under LMS conditions.

### Phenotypic and functional decrease in insulin signaling cascade

Excess free fatty acids have been reported as lipotoxic to hepatocytes and as initiators of inflammation processes that disrupt the glucose metabolism balance between glycolysis and gluconeogenesis,^53,55^ regulated by the insulin signaling cascade.^34,56–58^ Insulin receptor substrate 2 (IRS-2), a key membrane protein of the insulin receptor complex, has been reported to be more prone to proteolysis during high-fat diet events.^59,60^ Previously we have reported a significant decrease in the total IRS-2 signal in the vLAMPS under EMS.^1^ In the present study the decrease in signal was also significant under LMS conditions compared to NF and comparable to EMS (Figure 3 A to C). The average IRS-2 signal across the overall vLAMPS was statistically lower in both EMS and LMS compared to NF (Figure 3 C), with no significant signal difference between zones. The disruption of the insulin cascade mechanism was functionally indicated in the progressive decrease in insulin receptor-mediated insulin clearance under both EMS and LMS conditions (Figure 3 D), with significantly less clearance under LMS conditions starting at day 4 (p values in Figure 3 D). These studies demonstrate an association of one aspect of hepatic insulin resistance with increased steatosis in vLAMPS. Therefore, we sought to associate other aspects of insulin resistance, such as dysregulated gluconeogenesis with steatosis.

**Figure 3.**
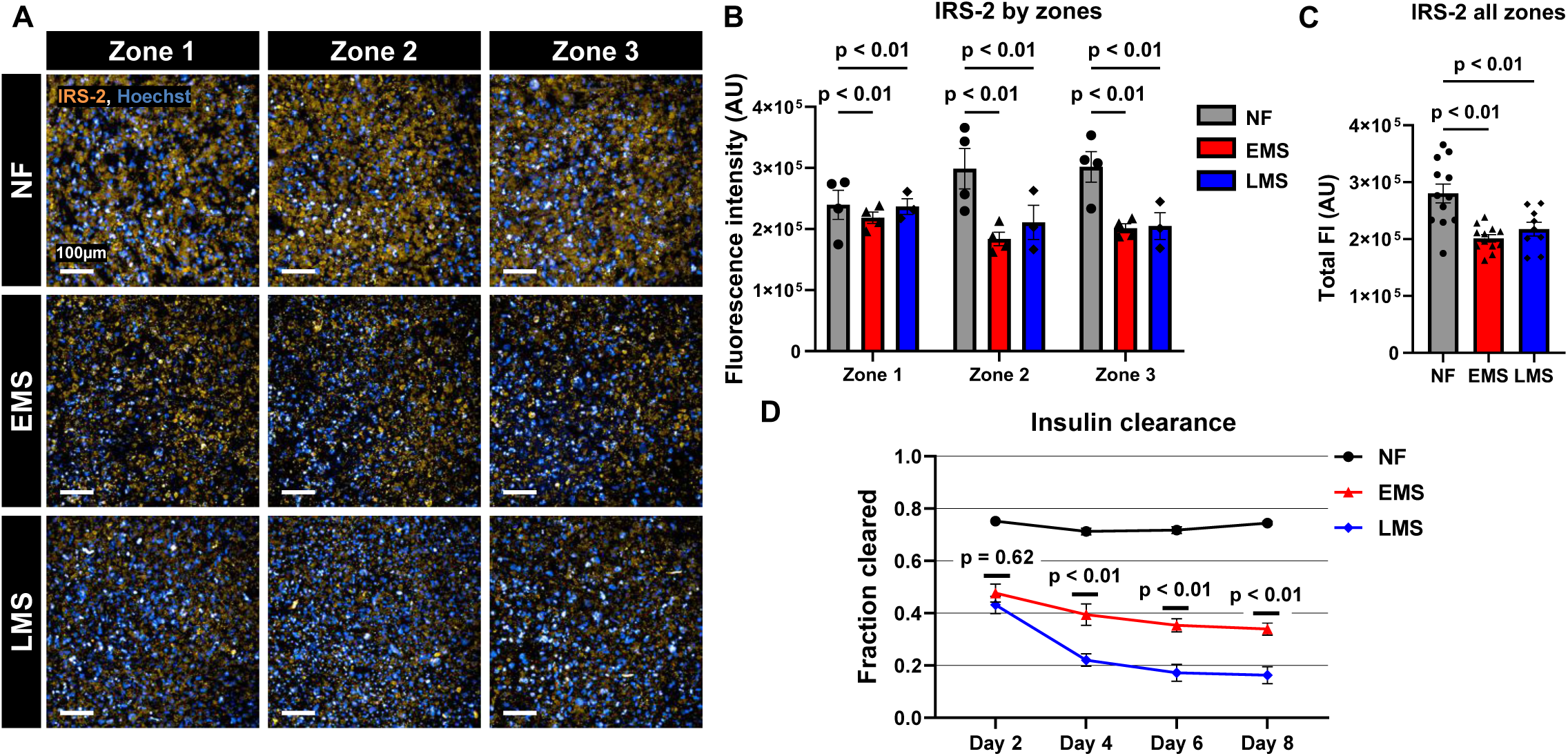
In vLAMPS the disease state reduces protein expression of Insulin receptor substrate 2 (IRS-2) and receptor mediated insulin clearance. (**A**) Representative immunofluorescence images of lipid droplets (LipidTOX) and IRS-2 in hepatocytes under NF, EMS, LMS conditions in the three zones. Scale=100µm. (**B**) IRS-2 signal quantification by zones. (**C**) The averaged quantification of IRS-2 in vLAMSP exhibits significant absence of signal in EMS and LMS medias. Scatter plot graphs display the mean and standard error of mean (SEM) of the fluorescence intensity of the IRS-2 signal of 8 fields at the ROI of each metabolic zone in the vLAMPS, n = 6 MPS. For graph -B-statistical analysis by Two-Way ANOVA with multiple comparison by Fisher’s LSD test to individually compare media condition with their respective zone with p-values shown. For graph -C-statistical analysis by One-Way ANOVA with multiple comparison by Dunnett test to compare EMS and LMS to NF medium with p-values shown. (**D**) Insulin clearance measured by fraction cleared between the detected insulin in the hepatic chamber efflux and their corresponding media influx. Line graphs display the mean and standard error of mean (SEM), n = 12 MPS. For graph -D-statistical analysis by Two-Way ANOVA with multiple comparison by Tukey test to compare the mean of each medium per day with p-values shown for EMS vs LMS.

### Insulin resistance manifested by dysregulated gluconeogenesis

Monitoring efflux glucose levels in reference to the glucose levels in the influx indicates both functional glucose metabolism and hepatic glucose production.^61,62^ The glucose consumption ratio (i.e., efflux/influx) from the hepatic chamber was constant across all three conditions under flow on day 2 (Figure 4 A). Under NF conditions, a steady decrease in this ratio was observed, indicating balanced glucose homeostasis with net glucose consumption in the vLAMPS. The glucose levels in the efflux of the EMS and LMS conditions increased. On day 8 the glucose efflux/influx ratio surpassed 1 under LMS conditions (Figure 4 A). These results suggest that in vLAMPS under late metabolic syndrome conditions glucose uptake is diminished and gluconeogenesis is increased, both classical indicators of hyperglycemia that are associated with hepatic insulin resistance.

**Figure 4.**
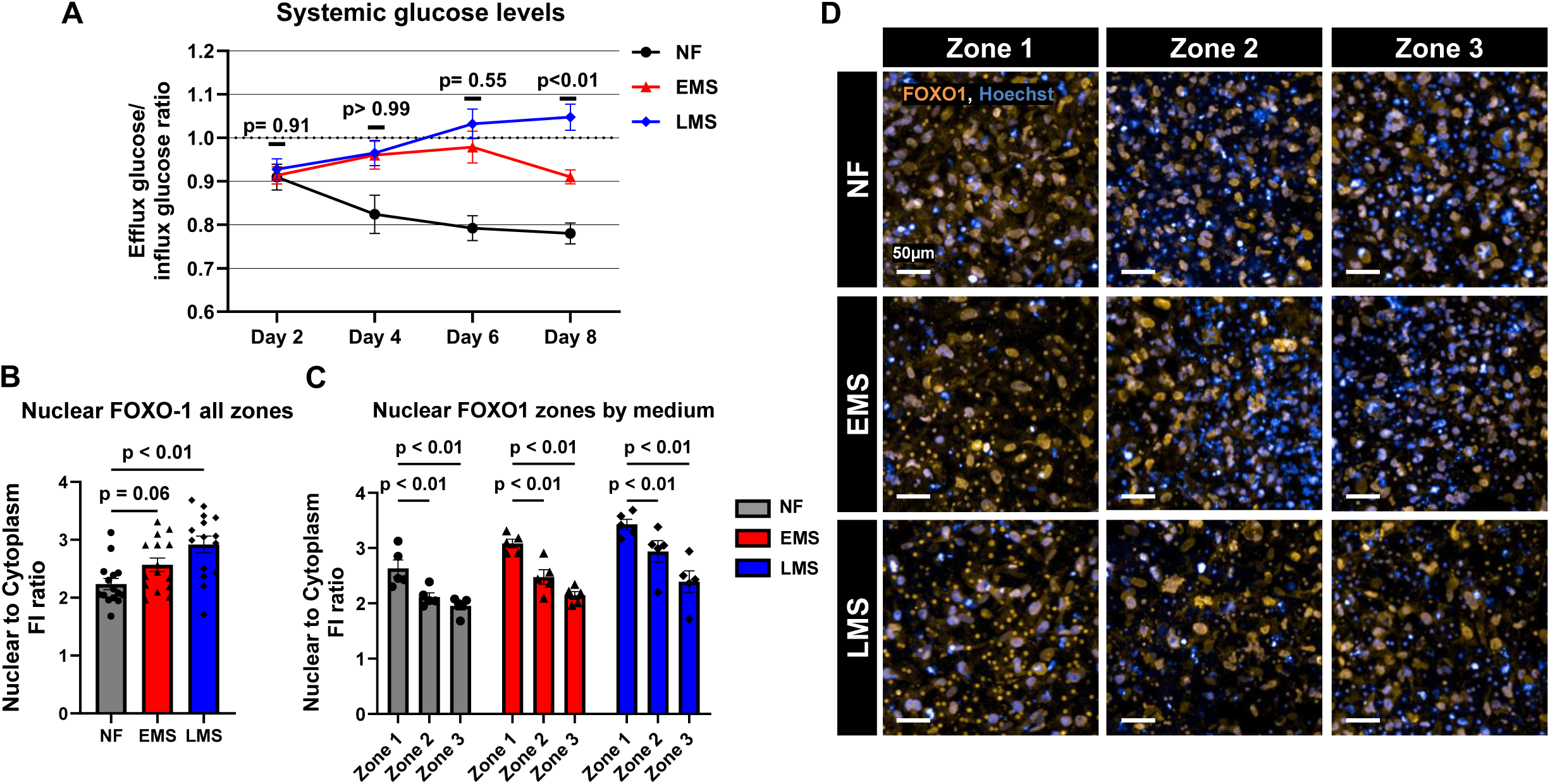
Disease severity increases systemic glucose production and hepatocyte specific nuclear translocation of FOXO1. (**A**) Ratio between the detected glucose in the hepatic chamber efflux and their corresponding media influx, with the LMS efflux having a glucose surplus over the influx. Line graphs display the mean and standard error of mean (SEM), n = 12 MPS. For graph -A-statistical analysis by Two-Way ANOVA with multiple comparison by Tukey test to compare the mean of each medium per day with p-values shown for EMS vs LMS. (**B**) The average nuclear to cytoplasmic ratio of the fluorescence intensity of all FOXO1 positive hepatocyte signal is significantly higher in EMS and LMS. (**C**) Nuclear to cytoplasmic ratio of the fluorescence intensity quantification of FOXO1 signal on CK-8 positive cells by zones under NF, EMS, or LMS perfusion on day 8. Scatter plot graphs display the mean and standard error of mean (SEM) of the fluorescence intensity of the nuclear FOXO1 signal of 8 fields at the ROI of each metabolic zone, n = 6 MPS. For graph -B-statistical analysis by Kruskal-Wallis with multiple comparisons by Dunn’s test to compare EMS and LMS to NF medium, with p-values shown. For graph -C-statistical analysis was performed using One-Way ANOVA and Dunnett multiple comparison test of the mean nuclear-to-cytoplasm ratio of each zone per medium condition, with p-values shown. (**D**) In their respective zones, representative immunofluorescence images of the hepatocytes (CK-8) and FOXO1 signal under NF, EMS, and LMS conditions. Scale=50µm.

We next investigated the molecular basis for enhanced glucose production by monitoring FOXO1 nuclear translocation. FOXO1 is a transcription factor that promotes gluconeogenesis. Its nuclear translocation is inhibited by insulin and is relatively increased under insulin resistance conditions.^21,63,64^ An elevated nuclear FOXO1 signal in comparison to its cytoplasmic signal across the sum in all zones was observed under LMS conditions, with an increasing trend under EMS compared to NF conditions (Figure 4 B). This result is consistent with the increase in glucose production inferred under both EMS and LMS conditions (Figure 4 A). We next determined the nuclear to cytoplasmic FOXO1 signal in each of the three zones across the vLAMPS. Under NF conditions, the nuclear to cytoplasmic signal of FOXO1 is significantly higher in Zone 1 relative to Zone 3 (Figure 4 C and D). This is consistent with higher gluconeogenesis known to occur in the periportal region in comparison to higher glycolysis known to occur in the oxygen poor pericentral zone.^52,55^ Although the nuclear to cytoplasmic FOXO1 signal was increased under both EMS and LMS conditions, this same zone-dependent (periportal > pericentral nuclear FOXO1) pattern was observed on day 8 (Figure 4 C). These results indicate that vLAMPS can recapitulate key structure and function aspects of the liver acinus under both normal and insulin resistant conditions.

### The role of metabolic and molecular drivers in the onset of acinar fibrosis

The results of these studies indicate in vLAMPS a significant association between hallmarks of insulin resistance and steatosis that are most evident under LMS conditions. In light of evidence suggesting a causal role of lipid-induced hepatic insulin resistance in MASLD progression,^65^ we also studied how NF, EMS, and LMS conditions affect later stage markers of MASLD. We focused on stellate cell activation and especially markers of fibrosis since these are most tightly associated with MASLD progression and its pharmacologically induced resolution. We observed that α-SMA, an *in situ* marker of stellate cell activation, was most significantly present under LMS conditions (Figure 5 A-C) with a trend towards a zone 3 preference. Correspondingly, the secreted markers of fibrosis and matrix remodeling, Col1A1 and TIMP1 were most significantly elevated under LMS conditions (Figure 5 D and E). Collectively, our results demonstrate a significant association between insulin resistance and both early and late stage MASLD progression in vLAMPS.

**Figure 5.**
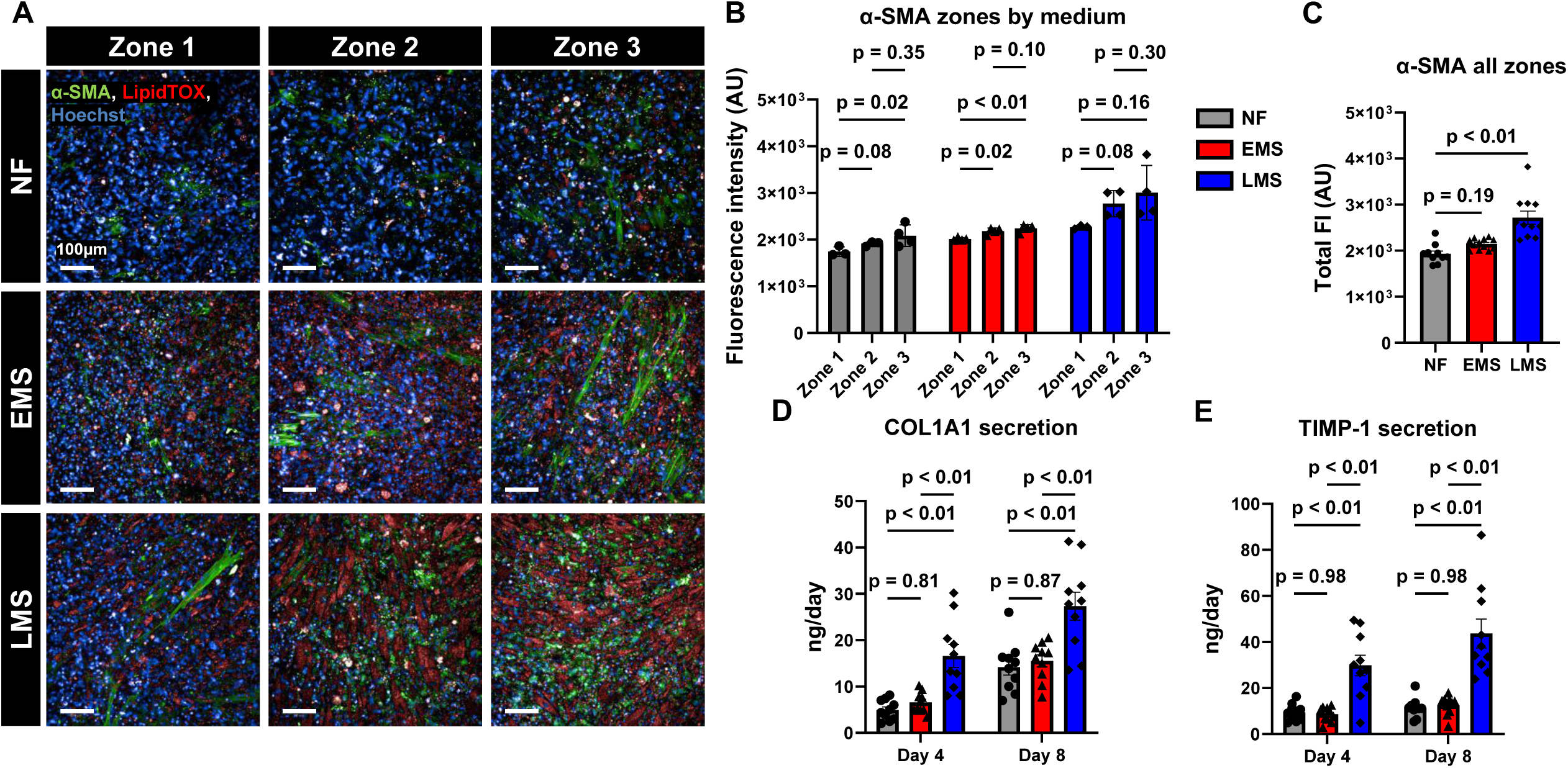
Late metabolic syndrome media induces stellate cell activation indicated by increases in α-SMA, collagen 1A1, and TIMP-1 expression. (**A**) Representative immunofluorescence images of lipid dropplets (LipidTOX) and α-SMA signal under NF, EMS, and LMS conditions in their respective zones. Scale=100µm. (**B**) Fluorescence intensity quantification of the α-SMA signal by zones under NF, EMS, or LMS perfusion on day 8. (**C**) The average fluorescence intensity of all α-SMA signal is significantly higher in LMS compared to EMS and NF. Scatter plot graphs display the mean and standard error of mean (SEM) of the fluorescence intensity of the α-SMA signal of 8 fields at the ROI of each metabolic zone in the vLAMPS, n = 5 MPS. For graph -B-, statistical analysis by Two-Way ANOVA with multiple comparison by Fisher’s LSD test to individually compare media condition with their respective zone with p-values shown. For graph -C-, statistical analysis was performed using one-way ANOVA with multiple comparisons using the Dunnett test to compare EMS and LMS to NF medium with p-values shown. The efflux levels from the hepatic chamber for (**D**) COL1A1 and (**E**) TIMP-1 are significantly higher in LMS than NF and EMS. Scatter plot graphs display the mean and standard error of mean (SEM), n = 12 MPS. For graphs -D and E-, statistical analysis was performed using two-way ANOVA with multiple comparisons using the Tukey test to compare the mean of each medium on their respective days, with p-values shown.

### Reproducibility of diverse functional and phenotypic measurements

The reproducibility of the above readouts is essential for establishing confidence in the measurements, thereby enabling each stage of MASLD to be defined and the underlying pathophysiology to be determined. We employed the Pittsburgh Reproducibility Protocol (PReP)^66^ to assess the inter-study reproducibility of the metrics. Albumin, Urea, lactate dehydrogenase (LDH), glucose level, and insulin clearance readouts in the hepatic and vascular chamber demonstrated significant reproducibility in amplitude and trend (interclass correlation coefficient, ICC) over the 8-day experiment across all three media, as detailed in Supplementary Table 2. Similarly, the *in situ* lipid signal intensity (LipidTOX), IRS-2 and FOXO1 signal, and VLDL readouts exhibited significant reproducibility in amplitude (coefficient of variation percentage analysis) at day 8 in all three media (Supplementary Tables 3). Similarly, the fibrotic markers associated with activated stellate cells (α-SMA) and fibrosis-related macromolecules COL1A1 and TIMP1^67–69^ were reproducible in trend and amplitude across independent studies (Supplementary Tables 2 and 3). These results support the utility of vLAMPS for comparative analyses and strengthen the validity of findings across diverse functional and phenotypical settings.

## Discussion

In this study, several functional and structural hallmarks of lipid-induced hepatic insulin resistance were observed in vLAMPS that were associated with MASLD progression. Given the significant pathophysiological role of hepatic insulin resistance in MASLD,^19^ these results are consistent with this human microphysiological system recapitulating MASLD progression through mechanisms identified in clinical studies and in genetically engineered and diet-induced *in vivo* models. Insulin receptor-mediated insulin clearance was significantly diminished as MASLD progressed from an early to late stage (Figure 3 C and D). This is consistent with prolonged exposure to elevated insulin levels and hepatocyte stress inducers dysregulating insulin receptor endocytosis, recycling, and lysosomal-mediated proteolytic degradation.^65,70^ Insulin receptor substrate-2 (IRS-2) protein expression was also significantly diminished. IRS-2 is known to selectively downregulate gluconeogenesis in comparison to lipogenesis^59,60,70^ and its reduced expression provides at least one mechanism for selective hepatic insulin resistance and the overproduction of hepatocyte-derived glucose observed in vLAMPS, a fundamental characteristic of MASLD progression. FFA influx into hepatocytes can also lead to uncontrolled upregulation of gluconeogenesis through FFA metabolic conversion to ceramides that inhibit the kinase activity of AKT, a critical downstream node of insulin receptor signaling.^19,20^ This selective inhibition of AKT prevents phosphorylation of the transcription factor, FOXO-1 enabling its nuclear translocation and resultant transcriptional activity to induce dysregulated gluconeogenesis.^63,71^ Enhanced nuclear translocation of FOXO-1 was observed in vLAMPS with increased MASLD severity (Figure 4 B). The nuclear translocation of FOXO-1 occurred selectively in zone 1, the periportal zone (Figure 4 C-D), the known primary site of hepatic gluconeogenesis. Increased flux of FFAs and their re-esterification to DAGs can activate PKCε to subsequently inhibit insulin receptor kinase activity by phosphorylating the insulin receptor at threonine 1160.^72^ This upstream inhibition of insulin receptor signaling will not only lead to enhanced hepatic gluconeogenesis but downregulation of the classical insulin pathway, INRSR-IRS-PI3K-AKT-mTORC1-SREBP-1c that normally results in *de novo* lipogenesis,^24,73,74^ thereby converting carbohydrates into storable triacylglycerols and other lipids. Despite the aforementioned indicators that insulin receptor signaling was increasingly compromised with increasing MASLD progression in vLAMPS, lipid droplets accumulated dyshomeostatically in the hepatocytes, preferentially in zone 3 (i.e., pericentral) of vLAMPS, recapitulating a key clinical phenotype of MASLD (Figure 2 A and B) that suggests alternative pathogenic *de novo* lipogenesis pathways in the insulin resistant state (see below). Similarly, VLDL secretion, normally limited by insulin-dependent activation of PI3K,^54,75,76^ increased with disease progression in vLAMPS (Figure 2 D), providing another indicator that vLAMPS can recapitulate the strong association of the hepatic insulin resistant state with MASLD progression.

Intrinsic to the complex pathophysiology of MASLD development are several extrahepatic factors that involve inter-organ crosstalk^77,78^ To account for critical aspects of these factors, we have modified the composition of the influx medium to mirror the metabolic syndrome state that clinically is closely associated with MASLD.^42,43^ Thus, both our early and late metabolic syndrome media contain FFAs to reflect the dysregulated lipolysis and diminished storage capacity in adipose tissue that causes overflow of FFAs to the liver. Similarly, in comparison to the normal fasting medium, the metabolic syndrome media contain elevated glucose to reflect in part the reduced capacity of skeletal muscle to take up glucose and store it as glycogen and the increased hepatic glucose output. Accordingly, the metabolic syndrome media also contain elevated levels of insulin mirroring the hyperinsulinemia that results from pancreatic beta cell glucose stimulated insulin secretion.^79^ To enable the development of chronic later stage disease that possesses inflammatory and fibrotic components without the need for prolonged incubation periods that could lead to confounding nonspecific cell viability issues, TGF-β and endotoxin (reflecting gut microbiome perturbation) were added to the late metabolic syndrome medium.^25,80,81^ Importantly, although donor cells originate from patients at different clinical stages, these standardized media formulations drive the uniform progression of MASLD *in vitro*, enabling controlled cross-donor comparisons while preserving genotype-specific biology and heterogeneous treatment responses.

These results support the use of vLAMPS for determining the underlying clinically relevant pathophysiology unique to the insulin resistant state and identifying targetable mechanisms involved in dysregulated *de novo* lipogenesis that drive MASLD progression. For example, in the hepatic insulin resistant state with high glucose influx and gluconeogenesis, the predominant pathway mediating carbohydrate-driven *de novo* lipogenesis involves the transcription factor, ChREBP, the carbohydrate response element-binding protein.^82^ ChREBP senses and is activated by glucose-6-phosphate, generated by glucokinase to initiate the transcription of a set of lipogenic genes that overlap with those more classically controlled by SREBP-1c in the normal state. Interestingly, a genome-wide association study (GWAS) has identified loss of function variants of glucokinase regulatory protein, GKBP, that controls cytoplasmic glucokinase enzymatic activity through regulating glucokinase cytoplasmic-nuclear translocation.^83^ These variants are associated with increased *de novo* hepatic lipogenesis and are a risk factor for MASLD.^84^ We suggest that vLAMPS can be used to identify a small molecule from a set of previously identified structurally diverse glucokinase binders that can mimic the critical modulatory properties of GKBP to modify dysregulated *de novo* lipogenesis and MASLD progression selectively in the insulin resistant state.

Recently two additional pathways involving the transcription cofactor, CREBZF,^85^ and the protein modulator, WDR6^86^ have been identified primarily from studies in mice that can upregulate *de novo* lipogenesis selectively in the insulin resistant state. vLAMPS can be used to corroborate and further characterize these pathways in the human setting as well as identify additional novel and targetable pathways enhancing *de novo* lipogenesis selectively in the insulin resistant state. We anticipate that these pathways will engender heterogeneity among patients in the insulin resistant state that is intrinsic to MASLD. MASLD heterogeneity arises from heritability as genome-wide association studies have identified both high and low risk genotypes, co-morbidities that as discussed above comprise the CKM syndrome, lifestyle decisions that involve weight and dietary control, and a perturbed gut microbiome that can disrupt the gut-liver axis promoting hepatic metabolic dysregulation and inflammation. With the conditional approval of the thyroid beta receptor agonist, resmetirom, and the emergence of a promising mechanistically diverse pipeline,^3^ there is an unmet need to understand how heterogeneity impacts MASLD pathophysiology, prognosis, and drug responses in individual patients. We expect that advancing the use of patient-specific differentiated iPSC-derived cells in vLAMPS (patient biomimetic twins) and its integration with computational approaches involving patient digital twins^38^ will help address the fundamental issue of heterogeneity for precision medicine. Finally, vLAMPS can be physically coupled with other organ-specific microphysiological systems^1^ to construct and deconstruct this holistic modular system to identify targetable mechanisms involving inter-organ crosstalk that could complement the developing pipeline of therapeutic strategies.

## Methods

### Human Hepatocytes and Non-Parenchymal Cells (NPCs)

Following our previously reported vLAMPS configuration,^1,35^ four key liver cell types (primary hepatocytes and liver sinusoidal endothelial cells) and 2 immortalized human cell lines (macrophages and human stellate cells) were sequentially seeded into the middle layer membrane of the vLAMPS. Cryopreserved human primary hepatocytes were secured from Thermo Fisher (HU8391, 62 years old male Caucasian with BMI of 22.2) and primary liver sinusoidal endothelial cells (LSECs) were purchased from LifeNet Health (HC170061, 79 years old female Africa American with BMI 27.8). Human hepatic stellate cell line LX-2 were acquired from Sigma (cat. no. SCC064) and phorbol myristate acetate (PMA) (Calbio, cat. no. 524400) treated THP-1 cells (ATCC cat. no. TIB-202) were used as Kupffer-like cells.

### vLAMPS Assembly

The assembly of the vLAMPS was carried out as previously described.^1,35^ Four days before initiating flow, the glass middle layer with a 77mm^2^ oval area and a polyethylene terephthalate (PET) membrane with 0.4µm pores (Micronit, cat. no. 03237) was coated with an ECM solution containing 100 μg ml^-1^ fibronectin (Sigma, cat. no. F1141) and 150 μg ml^-1^ collagen (Corning, cat. no. 354249) in 1x phosphate buffered saline (PBS) (Gibco, cat. no. 21600-069), and incubated overnight at 4°C. All non-endothelial cells were seeded based on the proportions of liver cell types *in vivo* determined by allometric scaling to be 40: 14: 6, Hepatocytes: Kupffer: Stellate.^35,87–89^ LSEC seeding ratio is 1:2.5 to hepatocytes to cover the surface area of the PET membrane as previously described.^1,35^ On the third day prior to flow, a solution of passage 2 LSECs in EGM-2 medium we seeded at a density of 8.5×10^5^ cells ml^-1^ on the vascular chamber (basolateral) side of the membrane. After an overnight incubation, the middle layer is flipped, and the remaining NPCs were seeded on the apical side of the membrane inside the oval-well. A blend of LX-2 at 1.3×10^4^ cells ml^-1^ and PMA-treated THP-1 at 2.4×10^4^ cells ml^-1^ prepared in EGM-2 medium and incubated overnight. To achieve hepatocyte polarization, the cells were seeded between decellularized porcine liver ECM (LECM) (provided by the laboratory of Dr. Badylak and Dr. Hussey^90^) and a type 1 collagen hydrogel overlay.^39^ Hepatocytes were seeded at 2.12×10^6^ cells ml^-1^ in hepatocyte plating medium (HPM) and incubated overnight. HPM was composed of Williams E medium (Life Technologies, cat. no. A1217601), 5 % FBS (Fisher Scientific, NC9525043), 1 % Pen-Strep (Hyclone, cat. no. SV30010), 2% L-glutamine (Hyclone, cat. no. SH30034.01) and 1% sodium pyruvate (Fisher Scientific, 11360070). The same medium is used in the preparation of the LECM at 400 µg ml^-1^. The collagen overlay was prepared at 2.5 mg ml^-1^ using the medium corresponding to the experimental conditions, in normal fasting (NF) medium or either early or late metabolic syndrome (EMS or LMS) for disease conditions. Both LECM and collagen steps require a 30-minute incubation for proper fibrillogenesis. Lastly, the middle layer was fully submerged in NF, EMS, or LMS medium and incubated until flow set up. All incubation steps took place at 37 °C in 5% CO2.

The vLMAPS is assembled inside the Micronit 4-chip holder (Micronit, cat. no. 4515). First four bottom chamber glass layers (Micronit, cat. no. OOC 00739) are placed in the base of the holder, with the channel gasket facing upwards to create the vascular chamber. The middle layers are then placed with the oval-well facing upwards, ensuring the basolateral side is in contact with the bottom layer. The top layer is then set on top of the middle layer, aligning the top chamber gaskets with the glass perforations of the middle layer, while keeping the oval-well facing up to establish the hepatic chamber. In the appropriate inlet and outlet ports of the holder’s lid are inserted the plastic ferrules and tubing. The system was meticulously sealed, ensuring proper alignment of all ferrules, gaskets, and perforations. Finally, the system is perfused in a parallel double syringe configuration at 35 µl hr^-1^ (Figure 1).

### Normal fasting and metabolic syndrome media

The media composition and concentrations are described in supplementary table 1. The base formulation for normal fasting (NF) and early metabolic syndrome (EMS) media have been previously described.^1,42^ The late metabolic syndrome (LMS) medium has been updated from our previously described version^42^ with the following modifications.^43^ LMS consists of EMS medium with the addition of 1 µg ml^-1^ lipopolysaccharides (LPS) (Sigma, cat. no. L-2630) and 10 µg ml^-1^ transforming growth factors beta (TGF-β) (R&D Systems cat. no. 240-B).

### Efflux Collection

Efflux medium from the vLAMPS was collected in glass vials, which were changed every other day for 8 days after initiating flow. Samples were collected from the hepatic and vascular chambers (Figure 1). The samples collected were maintained at 4°C until the completion of the experiment. Efflux samples were then aliquoted in 500 µl samples and stored at -20 °C until analysis.

### Secretome Measurements

Albumin, urea, LDH, COL1A1, and TIMP1 were measured as previously described.^1^ The Albumin assays (Bethyl Laboratories cat. no. A80-129A, A80-129, and E101) were performed in a 1:10 and 1:1 efflux dilution for the hepatic and vascular chambers, respectively. COL1A1 (R&D Systems, cat. no. DY6220-05) was measured at a 1:250 and 1:100 efflux dilution for the hepatic and vascular chambers, respectively. TIMP1 (R&D Systems, cat. no. DTM100) was measured at a 1:100 and 1:50 efflux dilution for the hepatic and vascular chambers, respectively. Urea (Stanbio Laboratory, cat. no. SB-0580-250) and lactate dehydrogenase (Promega, cat. no. G1780) were measured with no efflux dilution. Controls were run in duplicate, and only sample values inside the standard curve range were used for analysis.

Glucose levels were measured using the Amplex Red Glucose assay (Thermo Fisher, cat. no. A22189) on 96-well clear flat bottom polystyrene plates (Corning, cat. no. 3593) using a 1:500 dilution for all three media conditions and chambers, following the manufacturer’s protocol.

Insulin concentrations for the NF samples were carried out separately from the EMS and LMS samples due to the large detection range (1000-fold). Insulin in NF experiments was determined using the Alpco Ltd Ultrasensitive Insulin ELISA kit (Thermo Fisher, cat. no. 50-751-3655) following the manufacturer’s protocol with no dilution in the samples. The less sensitive version of the same kit, Alpco Ltd Insulin ELISA (Thermo Fisher, cat. no. NC0038324), was implemented for both EMS and LMS samples using a 1:20 dilution for both hepatic and vascular chambers.

Using the Cholesterol Assay kit – HDL and LDL/VLDL (Abcam, cat. no. ab65390), we determined levels of very low-density lipoprotein (VLDL) in NF, EMS, and LMS samples following the fluorometric readout protocol from the manufacturer on 96-well clear flat bottom black polystyrene plates (Corning, cat. no. 3603) with no dilution.

### vLAMPS Immunofluorescence Imaging

On day 8, the vLAMPS cells were fixed with 2% PFA at 4°C for 30 minutes. Following the previously described methodology,^1^ hepatocytes were immunofluorescence labeled for either cytokeratin 8 (CK-8) (1:200) (Invitrogen, cat. no. MA1-06318), insulin receptor substrate 2 (IRS-2) (1:250) (Abcam, cat. no. ab134101), or Forkhead box O1 (FOXO1) (1:150) (Sigma, cat. no. SAB3500507). Stellate cells were labeled for alpha-smooth muscle actin (α-SMA) (1:100) (Sigma, cat. no. A2547). Lipid droplets in the hepatocytes were stained with LipidTOX (1:500) (Invitrogen, cat. no. H34476) fluorescent staining (595/615nm). Cell nuclei were stained using Hoechst (1:2,000) (Thermo Fisher, cat. no. 62249).

To capture the appropriate metabolic zonation across the vLAMPS, images were taken at the specific fields in the previously determined^1^ ROI denoted *Zone 1*, *Zone 2*, and *Zone 3* across the vLAMPS oval area (Figure 1). The zone 1-periportal 2×4 grid of fields were captured between 3.9-5.2 mm from the edge of the oval-well close to the inlet, following the long axis of the oval. The zone 2-midlobular images were captured between 6.5-7.8 mm, and the zone 3-pericentral ones between 9.75-11.05 mm. Images were acquired with a Nikon 20× (0.45 NA) objective using the Operetta CLS High Content Scanning system (Perkin Elmer). Applying confocal mode and maximum projection acquisition mode, all images were taken with a 3.3µm stack size on a 160µm height, using the 490/525nm, 561/572, and 650/671nm excitation/emission settings. Fluorescence intensity analysis was performed using Fiji, following the methodology previously described.^1,35^ To exclude background fluorescence for LipidTOX, IRS-2, FOXO1, and α-SMA, the images were segmented with an intensity threshold based on a secondary antibody-only control. To measure the total fluorescence intensity (FI) of the lipid droplets (LipidTOX), particle quantification was applied by excluding small objects (<10 μm^2^). FOXO1 nuclear to cytoplasmic (N/C) fluorescence intensity ratio^91^ was determined only on cells positive for the CK-8 signal. Nuclear fluorescence intensity was determined by selecting nuclei positive only for CK-8 signal. A negative mask for the nuclei was generated from the initial screen and applied to the FOXO1 images, excluding any FOXO1 signal from outside the nuclei. Cytoplasmic fluorescence intensity was determined by selecting the remaining CK-8-positive cell area, minus the nuclei mask, and quantifying the FOXO1 signal in the “cytoplasm”. The fluorescence intensity for each zone is an average of 8 fields at the ROI of each metabolic zone. The average N/C ratio of all fields is the mean signal of all the metabolic zones per MPS.

### Reproducibility Analysis

Data from all studies were uploaded to the Quris AI, EveAnalytics™ platform. Inter-study reproducibility analysis followed the Pittsburgh Reproducibility Protocol (PReP)^66^ and used the Graphing/Reproducibility feature in the EveAnalytics^TM^ platform for statistical comparisons. For multi-timepoint metrics the interclass correlation coefficient (ICC) was used to assess the reproducibility of both the magnitude and trend of the response among replicates and studies. ICC values above 0.8 are considered to be excellent reproducibility and values between 0.2 and 0.8 are considered to be acceptable reproducibility. ICC below 0.2 indicate poor reproducibility. Coefficient of variation (CV) was used to analyze reproducibility for single time point metrics of functional and phenotypic readouts. The percentage value ranges the reproducibility status from Poor (CV≥15%), acceptable (5% < CV < 15%), and Excellent (CV ≤ 5%).^66,92,93^

### Statistical Analysis

Statistical comparisons between zones and media were made using PRISM (version 10.3.1). All data are presented as mean ± standard error (SEM) for n= 5–12 devices per study. Normality was first determined with all data by the Shapiro-Wilk test. When at least one of the groups or studies did not follow the Gaussian distribution, a nonparametric test was applied. A Two-Way ANOVA was used with column and row effect/interaction for zone and media comparison studies. Fisher’s LSD test carried out multiple comparisons between 6 families (3 media and 3 zones). One-way ANOVA with the Dunnett test was applied to studies with normal distribution for media only comparison. Kruskal-Wallis with Dunn’s test was applied for studies with no normal distribution. Multiple comparisons were carried out by comparing the average of each disease media (EMS or LMS) to NF. For comparisons between chambers and media, the One-Way ANOVA with Fisher’s LSD test was used. A comparison of the average value for each media condition for each day was carried out by Two-Way ANOVA. Multiple comparisons between 7 families (3 media and 4 days) were conducted using the Tukey test. For media per chamber per day, analysis, Two-Way ANOVA with Fisher’s LSD test was carried out between 8 families (6 media-chamber and 2 days).

## Data Availability

All of the source data and metadata pertaining to the findings presented in the main paper and supplementary information were uploaded in EveAnalytics (previously BioSystics-Analytics Platform and MPS-Database).^92^ vLAMPS studies ID numbers are: 1088, 1267, 1291, 1304, 1305, 1306, 1327, and 1328.

## Acknowledgement

The authors would like to acknowledge valuable discussions with members of the Organ Pathobiology and Therapeutics Institute (OPTIn) and the Pittsburgh Liver Research Center (PLRC). This project used shared instrumentation that was acquired with National Institute of Health (NIH) grant S10OD028450-NIH/OD (D.L. Taylor). We would like to acknowledge the following grants from the NIH: U24TR002632-NIH/NCATS (M.E. Schurdak and D.L. Taylor), UH3TR003289-NIH/NCATS (D.L. Taylor), UH3DK119973-NIH/NIDDK (D.L. Taylor), 5RO1DK135606-02 (M.T. Miedel), 4UH3TR004124-04 (M.T. Miedel), and 1U2CTR004863-NIH/NCATS (L. Vernetti, M.T. Miedel, M.E. Schurdak and D.L. Taylor).

## Author contributions

J.A. and L.V. conceptualized and designed the experiments under the supervision of A.M.S., M.E.S., M.T.M, V.K.Y., and D.L.T. J.A. performed all MPS experiments. J.A., L.V., R.D., and G.L. participated in the efflux and imaging quantification. J.A. and M.E.S. carried out the reproducibility analysis. J.A., A.M.S., M.T.M., and M.E.S. wrote the manuscript under the supervision of V. K. Y. and D.L.T. All authors have read and agreed to the published version of the manuscript.

---

**Supplementary Table 1.**
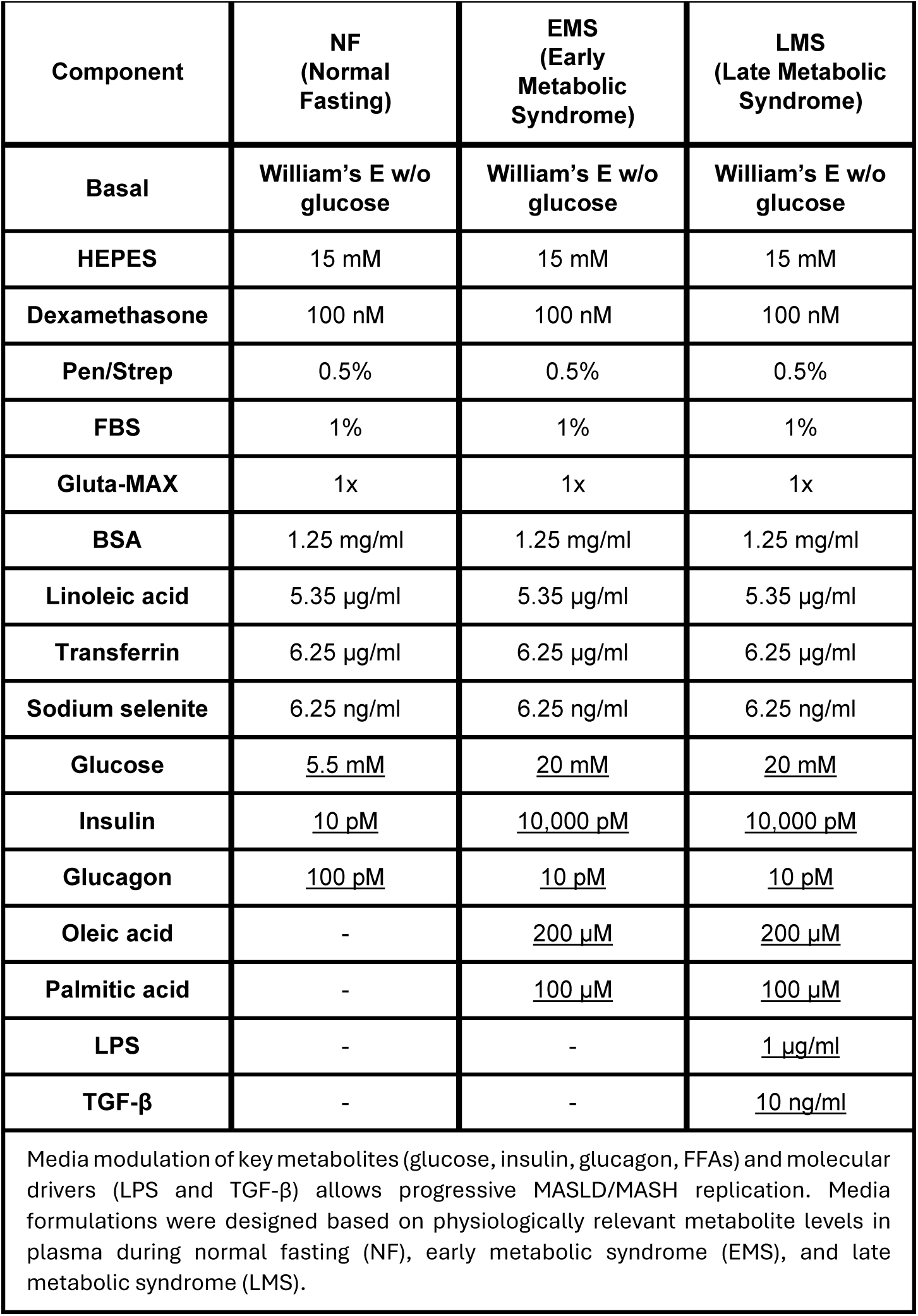
Media composition for the normal fasting, early metabolic syndrome, and late metabolic syndrome states used in the vLAMPS.

**Supplementary Figure 1.**
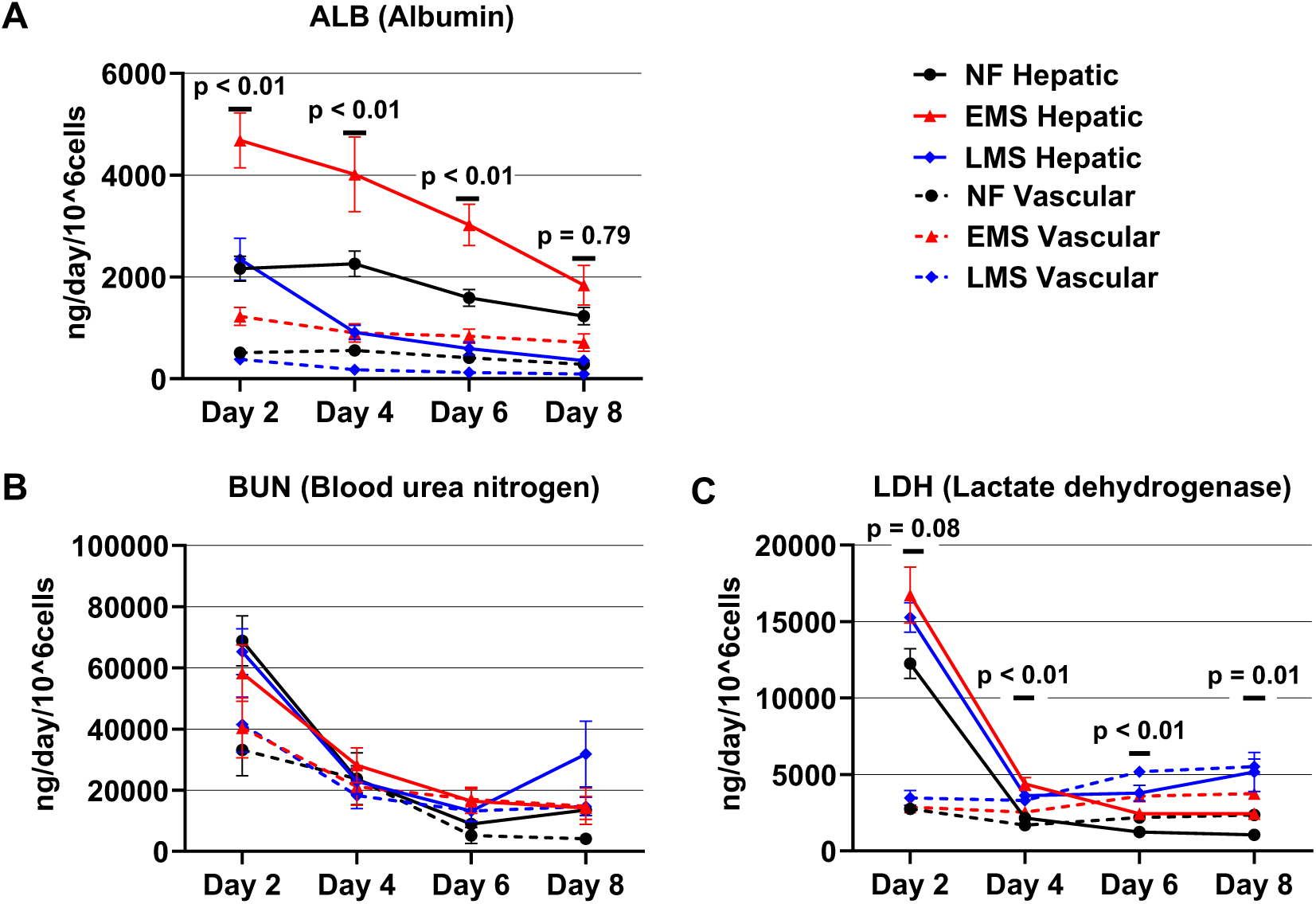
Basic liver functions were maintained in vLAMPS under NF, EMS, and LMS conditions. (**A**) The hepatic channel signal represents 81%±7, 82%±5, 80%±5, and 75%±10 of the total albumin values for days 2, 4, 6, and 8, respectively, in all three media. (**B**) Urea and (**C**) LDH secretions were consistent in both hepatic and vascular chambers across the 8 days. Line graphs display the mean and standard error of mean (SEM). For graph -A-Statistical analysis was done by Two-Way ANOVA and Sidak’s multiple comparison test to the mean albumin secretion between NF and EMS hepatic chamber values, with p-values shown. For graph -C-statistical analysis by Two-Way ANOVA with multiple comparison by Dunnett test to compare the mean LDH value in each day of EMS or LMS to NF medium from the hepatic chamber, with p-values shown.

**Supplementary Table 2.**
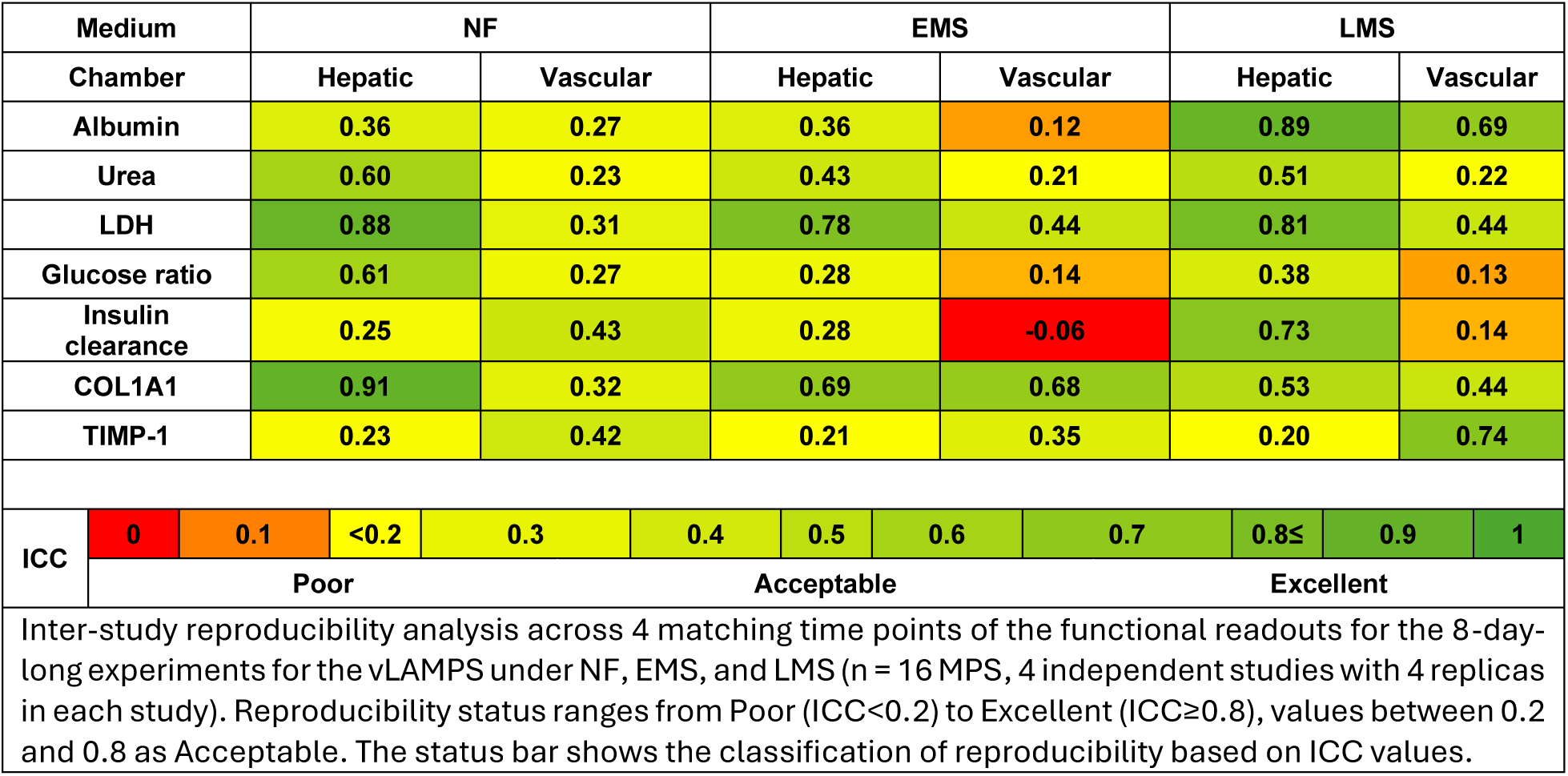
Interclass correlation coefficient (ICC) demonstrates an Acceptable reproducibility of both the amplitude and trend of the functional readouts in the vLAMPS in all three media through 8 days.

**Supplementary Table 3.** The reproducibility of functional and phenotypic analytes at a single time point on day 8 shows mixed trends ranging from Poor to Excellent by coefficient of variation (CV) analysis.

| Medium |  | NF |  |  |  | EMS |  |  | LMS |  |  |
| --- | --- | --- | --- | --- | --- | --- | --- | --- | --- | --- | --- |
| Chamber |  | Hepatic |  | Vascular |  | Hepatic |  | Vascular | Hepatic |  | Vascular |
| VLDL |  | 32.3 |  | 17.5 |  | 17.0 |  | 11.2 | 7.1 |  | 24.9 |
| LipidTOX |  | 10.2 |  |  |  | 3.3 |  |  | 7.5 |  |  |
| FOXO-1 |  | 7.9 |  |  |  | 18.4 |  |  | 14.7 |  |  |
| IRS-2 |  | 0.6 |  |  |  | 6.0 |  |  | 18.1 |  |  |
| α-SMA |  | 0.9 |  |  |  | 4.5 |  |  | 5.6 |  |  |
| CV | 0 | 2 | 4 | 6 | 8 | 10 | 12 | 14 | 16 | 18 | 20 |
|  | Excellent |  |  |  | Acceptable |  |  |  | Poor |  |  |
| Inter-study reproducibility analysis functional and phenotypic readouts using the CV percentage value for the vLAMPS under NF, EMS, and LMS on day 8 (n = 16 MPS, 4 independent studies with 4 replicas in each study). Reproducibility status ranges from Poor (CV≥15%), acceptable (5% < Max CV < 15%), and Excellent (CV ≤ 5%). The status bar shows the classification of reproducibility based on CV% values. The functional VLDL readout was carried out from outflow samples from the hepatic and vascular chambers. The phenotypic readouts (LipidTOX, FOXO-1, IRS-2, and α-SMA) represent the image z-stack analysis of the middle layer between the hepatic and vascular chamber. A Poor score can be indicative of a dynamic trend. |  |  |  |  |  |  |  |  |  |  |  |

